# Synaptotagmin isoforms differentially regulate glutamate and GABA release in the lateral habenula

**DOI:** 10.64898/2026.04.02.716068

**Authors:** Dustin N. White, J. Keenan Kushner, Kelly E. Winther, Dillon J. McGovern, Tamara Basta, Zoe R. Donaldson, Charles A. Hoeffer, David H. Root, Michael H. B. Stowell

**Affiliations:** Department of Molecular, Cellular & Developmental Biology, University of Colorado, Boulder, CO, USA; Institute for Behavioral Genetics, University of Colorado Boulder, CO, 80309 USA; Department of Integrative Physiology, University of Colorado Boulder, CO 80309, USA; Department of Psychology & Neuroscience, University of Colorado Boulder, Boulder, CO 80309, USA

## Abstract

Neurotransmitter co-transmission contributes to diverse physiological processes throughout the mammalian brain, including sensory integration, motivational control, and social behaviors. Projections from the globus pallidus internus (GPi; the entopeduncular nucleus, EPN, in rodents) to the lateral habenula (LHb) are well-characterized by the co-transmission of both GABA and glutamate. These dual-release inputs modulate behavioral states in chronically learned helpless (cLH) rats, influencing both the onset and recovery of pathological phenotypes. Here, we employed confocal 3D reconstructions that confirmed the presence of both vesicular transporters VGAT and VGLUT2 in EPN axon terminals within the LHb. Further investigation revealed that GABA and glutamate are packaged in distinct vesicle populations within individual presynaptic terminals. Notably, the calcium (Ca²⁺) sensors Synaptotagmin-2 (Syt2) and Synaptotagmin-3 (Syt3) are highly expressed in the EPN whereas expression of the canonical Ca²⁺ sensor, Synaptotagmin 1 (Syt1), is downregulated. Additionally, using confocal microscopy, we observed selective spatial correlations of Syt2 and VGLUT2 and between Syt3 and VGAT in LHb axon terminals. These observations strongly suggested that Syt2 serves as the predominant Ca²⁺ sensor for glutamatergic vesicle fusion, and Syt3 serves as the predominant Ca²⁺ sensor for GABAergic vesicle fusion in the LHb. To test this hypothesis, we employed targeted antisense oligonucleotide (ASO) knockdown of Syt2 and Syt3 in EPN neurons and measured LHb glutamatergic and GABAergic currents. Syt2 knockdown resulted in an increase in mEPSC frequency, amplitude, half-width and decay, suggesting increased glutamate vesicle release probability and increased glutamate vesicle packing. However, Syt2 knockdown had no influence on mIPSCs amplitude or frequency. On the other hand, Syt3 knockdown had no apparent effect on glutamate release but caused an increase in mIPSC frequency suggesting increased quantal release probability of GABA. Together, these findings identify a molecular mechanism by which synaptotagmin isoforms govern differential glutamate and GABA release at EPN dual-transmitter terminals in the LHb. These results provide evidence for presynaptic mechanisms regulating excitatory–inhibitory balance within this brain structure and importantly provide molecular targets for pharmacological intervention.

## INTRODUCTION

Henry Dale first postulated that each neuron releases a single neurotransmitter at its synapses, but accumulating evidence has overturned this seemingly universal rule^1^. Once thought nonexistent or anomalous, fast dual-release neurons have now been identified in major circuits throughout the mammalian brain^2–6^. In particular, glutamate and GABA, are co-released in EPN→LHb pathways that exhibit hyperexcitability in animal models with depressive phenotypes^7,8^. These neurotransmitters are often viewed dichotomously—glutamate as excitatory and GABA as inhibitory^9,10^. While their broad roles are generally consistent, the strict anatomical separation between glutamatergic and GABAergic neurons is less rigid than previously believed. Indeed, multiple examples demonstrate that glutamate and GABA can be synthesized, trafficked, and released at single axon terminals^11–13^ and more recently it has been proposed that glutamate and GABA are co-packaged within the LHb^14^.

We reasoned that if regulated release of opposing neurotransmitters does occur, it is likely due to isoform variations in synaptotagmin Ca²⁺ sensors^15^. Of the approximately seventeen known synaptotagmin isoforms^16^, several act as presynaptic Ca²⁺ sensors with distinct affinities and kinetic properties, with Syt1 being the most well established Ca^2+^ sensor in the mammalian brain^17–19^. We utilized the Allen Brain Atlas^20^ and determined both Syt2 and Syt3 transcripts are notably enriched in the presynaptic terminals of the EPN→LHb pathway compared to the thalamus, a region lacking dual-release properties. Furthermore, Syt1 transcripts were notably decreased in the EPN and the increase of Syt2 and Syt3 co-expression suggested a potential mechanism for regulating transmitter-specific vesicle fusion by segregating neurotransmitter transporters and Ca²⁺ sensors to individual synaptic vesicles. In alignment with this hypothesis, the two isoforms differ markedly in Ca^2+^ affinity, potentially staggering their responses to presynaptic Ca^2+^ influx—particularly if each associates with distinct vesicle populations^17^. Moreover, Syt2 mediates fast, synchronous vesicular release, whereas Syt3 has been linked to slower, asynchronous or modulatory functions^21,22^. Together, these observations led us to hypothesize that Syt2 and Syt3 could differentially regulate glutamate and GABA release at dual-transmitter terminals within the LHb, thereby tuning excitatory–inhibitory balance. To test this hypothesis, we designed a series of experiments that explored the colocalization of neurotransmitter transporters and synaptoptagmins in EPN→LHb projection terminals using confocal microscopy and the electrophysiological consequences of spatially targeted FANA-ASO knockdown of synaptoptagmin isoforms (**Figure 1)**.

**Figure 1.**
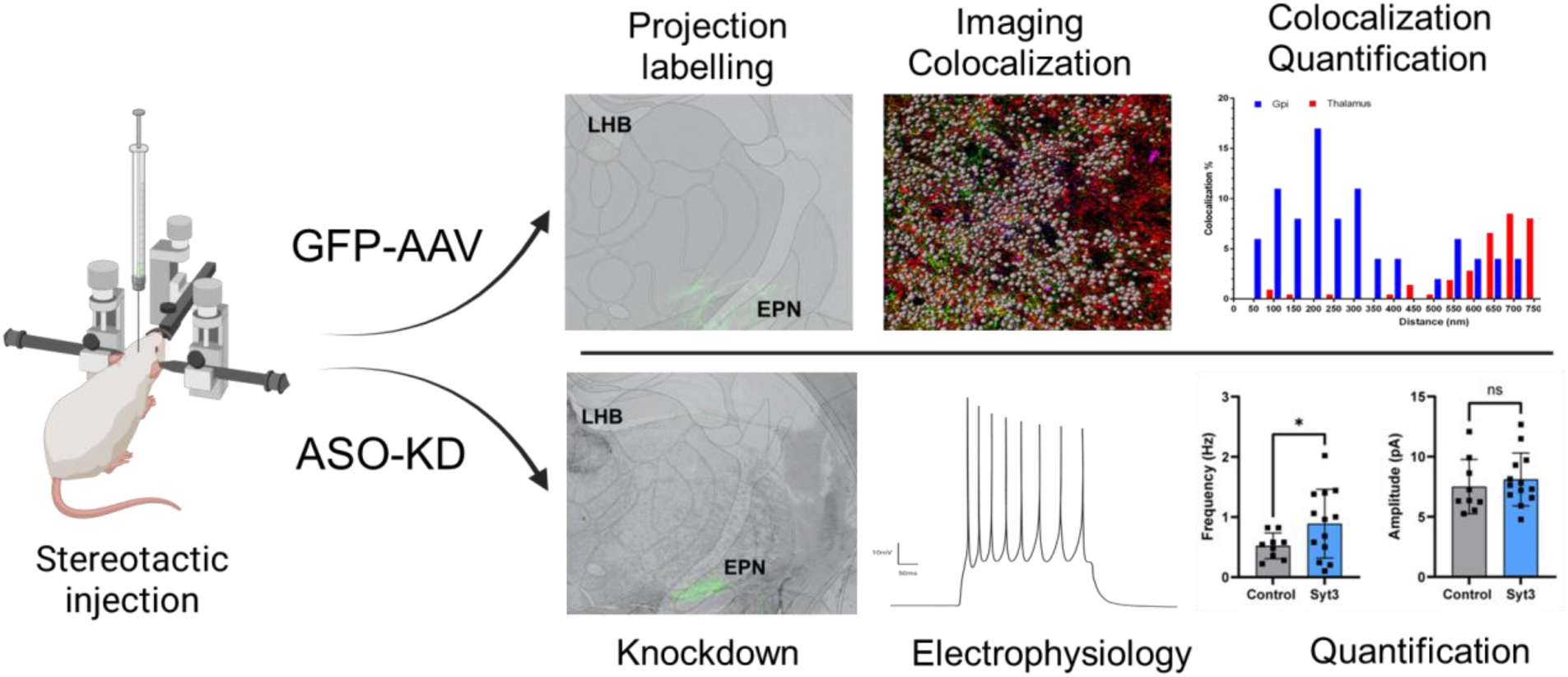
Overview of experimental workflow used to identify, quantify and knock down synaptotagmin isoforms in the LHB. Stereotactic injection of ASOs targeting a specific synaptotagmin or ChannelRhodopsin-AAV to mark projections from the EPN to the LHB were used to alter activity (top) or identify projections for colocalization (bottom).

## RESULTS

### VGLUT2 and VGAT are co-localized in EPN projections to the LHb

We performed immunolabeling and confocal imaging of vesicular transporters VGLUT2 and VGAT within the LHb. While both transporters were abundant in the LHb and the majority of VGLUT2 and VGAT puncta were in proximity, it was initially unclear whether they were confined to the same membrane-bound presynaptic bouton. To address this, an AAV encoding Cre-dependent CoChR-GFP was stereotactically injected into the EPN of VGluT2-IRES:Cre mice to fluorescently label the EPN→LHb projections. GFP-fluorescence was observed throughout the soma, axon, and synaptic boutons by high-resolution confocal imaging (**Figure 2B)**. Subsequent immunohistochemical analysis revealed that both VGLUT2 (glutamatergic) and VGAT (GABAergic) were co-localized within the labeled EPN→LHb projection terminals (**Figure 2C,D)**. Specifically, the spatial distribution of these transporters fell within the established threshold of focal error (0.1 micron) of the microscope for overlap, providing evidence for a dual-transmitter synapse in this specific circuit. In contrast, thalamic VGLUT2 and VGAT puncta not associated with EPN projections did not colocalize, consistent with the absence of dual-release activity in this brain region (**Figure 2 D,E)**. These observations are concordant with prior reports that observed colocalization of VGLUT2 and VGAT within the rat LHb^23^. This prior study also determined that isolated synaptic vesicles from the LHb have distinct populations of glutamate and GABA containing vesicles^23^. Are results demonstrate that these dual synapses primarily arise from EPN projections and are spatially localized within individual synaptic boutons.

**Figure 2.**
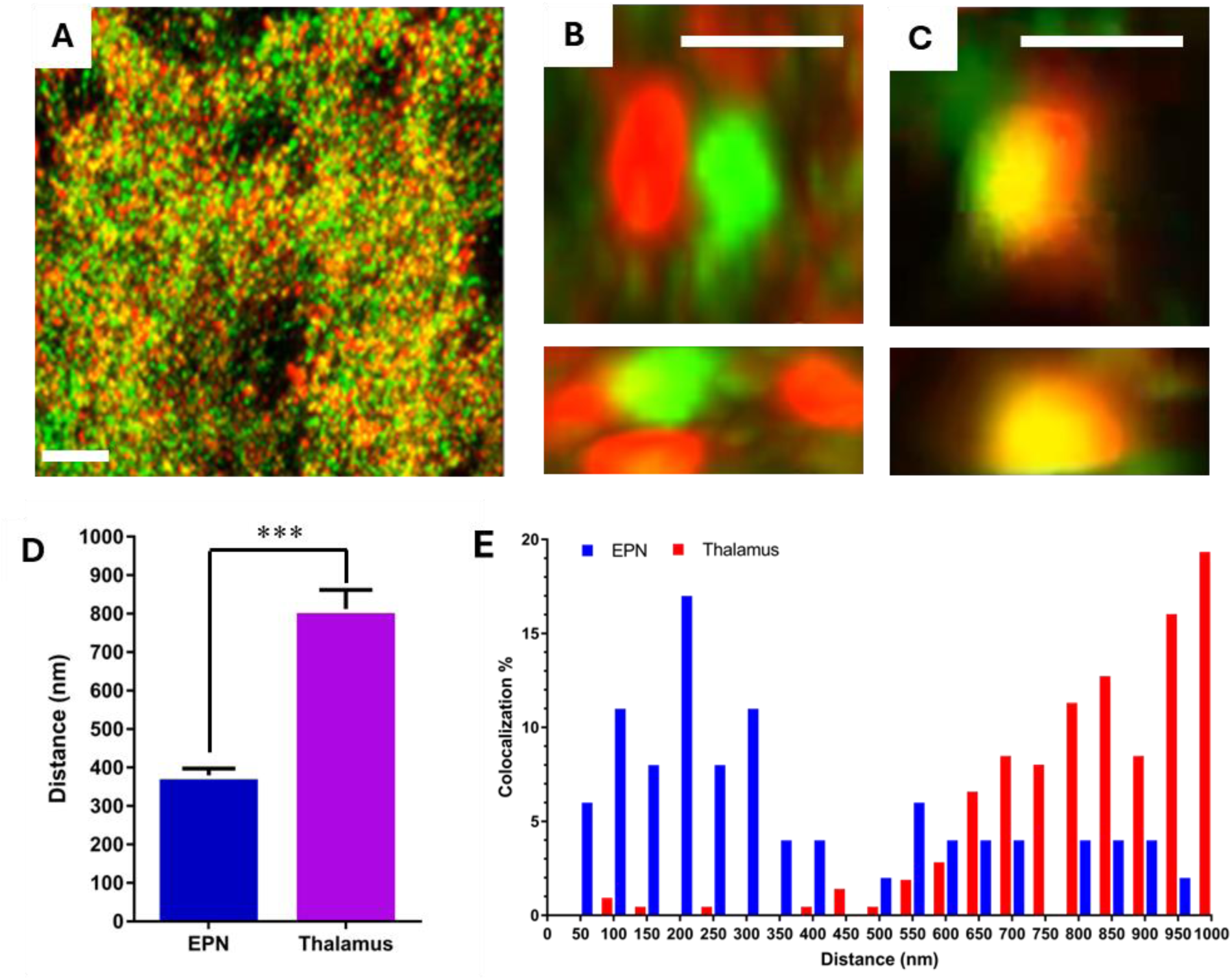
Colocalization of VGLUT2 and VGAT within EPN projections to the LHb. **A**) Confocal imaging of the LHb of EPN neurons shows distinct yet overlapping puncta for VGLUT2 (false color green) and VGAT (false color red). Both transporters localize within individual EPN→LHb terminals, indicating segregation of glutamate and GABA containing vesicles in the same bouton, scale bar 20 μm. **B,C**) Higher magnification of separate and co-labeled regions in XY(top) and YZ(bottom), scale bar 2 μm. **D**) Average distance between VGLUT2 and VGAT puncta in the LhB and thalamus respectively, n=3 mice total, *statistical comparisons by unpaired t-test. ***p < 0.001.* **E**) Distance distribution of VGLUT2 and VGAT colocalization observed in the EPN (blue) and thalamus (red).

### Syt2 and Syt3 transcripts are enriched in EPN neurons and colocalize to VGLUT and VGAT respectively

Having identified the EPN→LHb terminals as both glutamatergic and GABAergic, we sought to identify molecular mechanisms that could potentially regulate glutamate and GABA release independently. We utilized the genome wide in-situ hybridization (ISH) data from the Mouse Allen Brain Atlas^20^ to identify differentially expressed genes between the EPN and the thalamic reticular nucleus, a region presumed to be devoid of dual release synapses. We observed that the synaptotagmin transcripts Syt2 and Syt3 are markedly enriched in the EPN (**Figure 3A**). Interestingly, several other synaptotagmin isoforms transcripts, Syt1 in particular, were diminished in the EPN. This suggested that this enrichment is not a general family-wide phenomenon but rather specific to Syt2 and Syt3. Given the known differences in Ca^2+^ sensitivity and vesicle fusion kinetics^17,18^, we hypothesized that Syt2 and Syt3 could serve distinct functional roles within the EPN→LHb projections. Importantly, this specialization is consistent with models proposing temporal separation of vesicle fusion events as a mechanism of dual release^24^.

**Figure 3.**
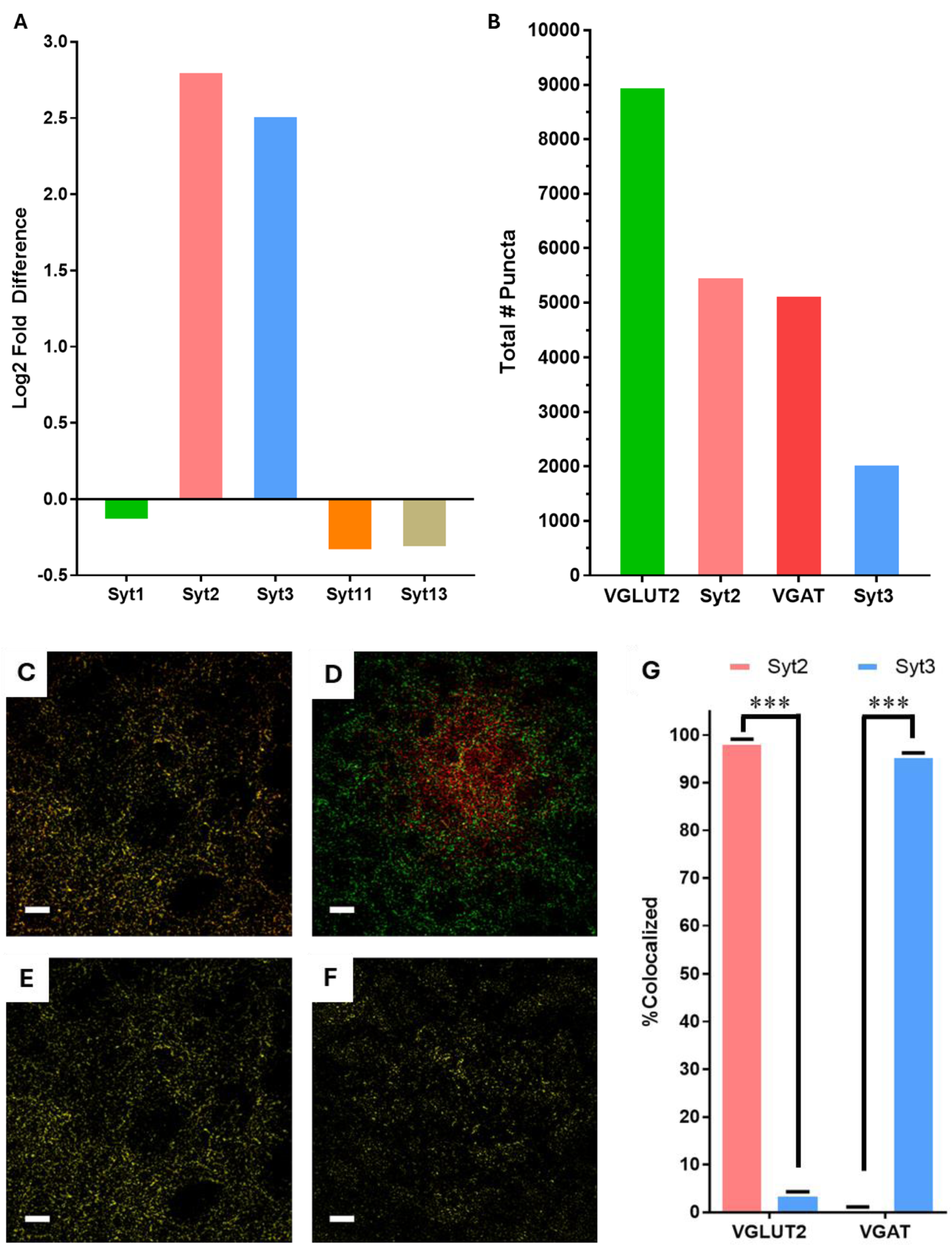
Syt2 and Syt3 are highly expressed and differentially associated with vesicular glutamate and GABA transporters within EPN axon terminals of the LHb. **A**) Differential expression levels of Syt1, Syt2, Syt3, Syt11, Syt13 between the LHb and Thalamus from the Allen Brain Atlas^20^. **B**) Exemplary confocal microscopy image showing col-localized Syt2(false color red) with VGLUT2(false color green) utilized for quantification in G, scale bar 10 μm. **C**) Colocalization image (yellow) calculated from the image in B, scale bar 10 μm. **D**) Exemplary confocal microscopy showing col-localized Syt3(false color red) with VGAT(false color green) utilized for quantification in G, scale bar 10 μm. **E**) Colocalization image (yellow) calculated from the image in D, scale bar 10 μm. **F**) Histogram of the total number of puncta analyzed for colocalization determination in G. **G**) Immunohistology overlap analysis of vesicle-type specificity of synaptotagmin isoform distribution. A total of 98% of the Syt2 signal overlapped with VGLUT2 puncta, whereas a total of 95% of Syt3 signal overlapped with VGAT, error bars SEM, n=3 mice total. *statistical comparisons by unpaired t-test. ***p < 0.001*.

We next examined whether each Syt isoform is preferentially associated with glutamate or GABA vesicle populations^17^. We reasoned that, under a dual-release rather than co-packaging model, transmitter-specific membrane protein composition could alter differences in vesicle fusion. Confocal imaging of over 10,000 EPN→LHb synaptic terminals (**Figure 3B**) combined with quantitative colocalization analysis revealed striking specificity (**Figure 3C-F**). Of all Syt2 puncta located within GFP-labeled EPN membranes, approximately 98% overlapped with VGLUT2. Conversely, 95% of total Syt3 signal was associated with VGAT rather than VGLUT2 (**Figure 3G**). These results demonstrate that within EPN→LHb terminals, Syt2 is primarily segregated on the glutamatergic (VGLUT2) vesicles and Syt3 is primarily segregated on GABAergic (VGAT) vesicles.

### Syt2 mediates excitatory transmission and Syt3 mediates inhibitory transmission

To assess the functional role of each isoform, we conducted spatially targeted antisense oligonucleotide (ASO) knockdown of Syt2 and Syt3. FAN-ASOs targeting either Syt2 or Syt3 were stereotactically injected into the EPN. Subsequently, we recorded miniature postsynaptic currents in the LHb and, following electrophysiological recording, knock down was assessed using western blot analysis of dissected tissue from both the treated and contralateral EPN Due to the cell type heterogeneity and imprecision of dissection, samples were normalized to total protein and Syt2 and Syt3 bands quantified to assess relative protein levels. This analysis was moderately informative and showed variable knockdown efficiencies (**Figure S2**). Nonetheless, knockdown of Syt2 produced significant alterations in several parameters associated with excitatory synaptic events, but not inhibitory events (**Figures 4**). Both the frequency and amplitude of miniature EPSCs were significantly increased following Syt2 knockdown indicating that Syt2 normally constrains glutamatergic vesicle release (**Figure 4**). These findings suggest that Syt2 serves as a key Ca²⁺ sensor governing fast, synchronous glutamate release at EPN→LHb terminals. Importantly, this trend was not observed for inhibitory events, as miniature IPSCs remained unchanged following EPN Syt2 knockdown. Thus, Syt2 appears regulate glutamatergic vesicle relase within these dual-release boutons. Using the identical procedure for Syt2, ASO-mediated knockdown of Syt3 had no effect on mEPSC but significantly increased the frequency of miniature IPSCs (**Figure 5).** This effect supports a complementary role for Syt3 as the principal Ca²⁺ sensor for GABAergic vesicle release in the EPN→LHb pathway. Together, these findings demonstrate that Syt2 and Syt3 functionally segregate to VGLUT2 and VGAT vesicle populations, respectively, conferring Syt isoform-specific control of glutamate and GABA release in the lateral habenula.

**Figure 4.**
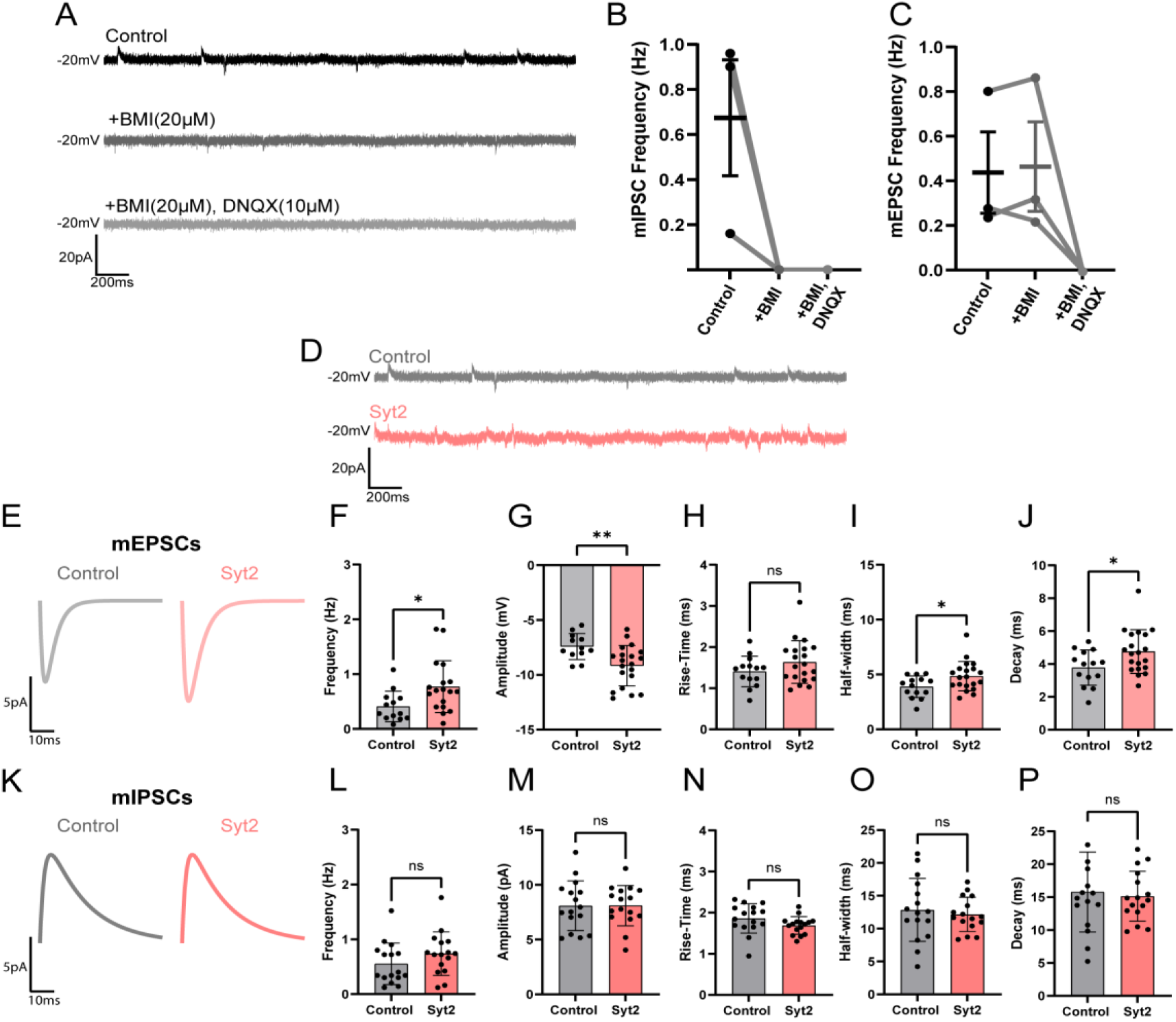
Syt2 selectively modulates excitatory but not inhibitory miniature synaptic transmission in LHb neurons. **(A–C)** Representative traces and quantification of miniature postsynaptic currents recorded from LHb neurons under control conditions and after application of the GABA_A_ receptor antagonist bicuculline methiodide (BMI, 20 μM) and AMPA receptor blocker DNQX (10 μM) (A). BMI abolished mIPSCs, confirming GABAergic identity (B), while DNQX had no effect on mIPSCs but blocked mEPSCs, validating pharmacological isolation of miniature excitatory and inhibitory currents (C). **(D–J)** Representative miniature recordings from control (gray) and Syt2-expressing (light red) LHb neurons (D). Average mEPSC from control and Syt2 ASO knockdown (E). Syt2 ASO knockdown significantly increased mEPSC frequency (F) and amplitude (G), with no change in rise time (H). Syt2 knockdown broadened mEPSC half-width (I) and prolonged decay time (J), indicating altered excitatory release kinetics and/or receptor activation. **(K–P)** Average mIPSC traces from control (gray) and Syt2 ASO knockdown (light red) neurons (K). Syt2 ASO knockdown did not significantly impact mIPSC frequency (L), amplitude (M), rise time (N), half-width (O), or decay (P), indicating that Syt2 selectively modulates excitatory (glutamatergic) but not inhibitory (GABAergic) transmission. *Data shown as mean ± SD; control n=16; Syt2 n=22; statistical comparisons by unpaired t-test. *p < 0.05, **p < 0.01; ns, not significant*.

**Figure 5.**
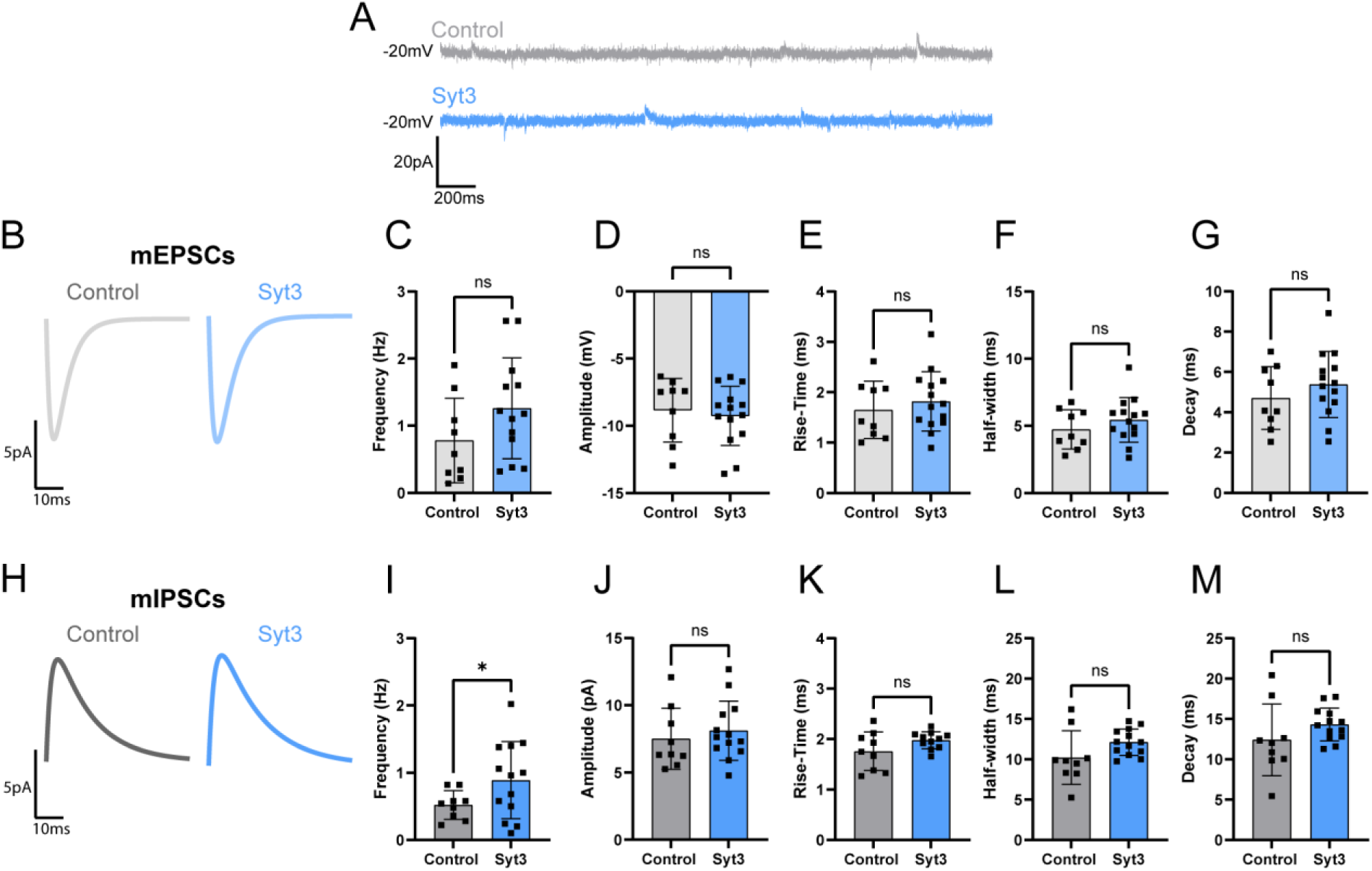
Syt3 knockdown selectively modulates inhibitory but not excitatory miniature synaptic transmission in LHb neurons. **(A–G)** Representative voltage-clamp traces (A) and summary analyses (B–G) of miniature excitatory postsynaptic currents (mEPSCs) recorded from control (gray) and Syt3 ASO-treated (light blue) LHb neurons. Knockdown of Syt3 did not significantly alter mEPSC frequency (C), amplitude (D), rise time (E), half-width (F), or decay kinetics (G), indicating that excitatory synaptic transmission is unaffected by Syt3 depletion. **(H–M)** Average mIPSC response (H) and quantification of miniature inhibitory postsynaptic currents (mIPSCs) recorded under control and Syt3 knockdown conditions. Syt3 ASO increased mIPSC frequency (I) without affecting event amplitude (J), rise time (K), half-width (L), or decay (M), consistent with a selective role for Syt3 in mediating spontaneous GABAergic vesicle release. *Data shown as mean ± SD; control n=10, Syt3 n=14; statistical significance determined by unpaired t-test. *p < 0.05; ns, not significant*.

## DISCUSSION

In this study we provide evidence for a molecular mechanism by which specific synaptotagmin isoforms govern differential neurotransmitter release in a dual-transmitter circuit. Using anatomical, molecular, and functional approaches, we demonstrate that projections from the globus pallidus internus (GPi; entopeduncular nucleus, EPN, in rodents) to the lateral habenula (LHb) contain distinct vesicle populations for the accumulation of glutamate (via VGLUT2) and GABA (via VGAT), each associated with a specific synaptotagmin isoform. Syt2 and Syt3 are both enriched in the EPN, but their localization and function diverge: Syt2 localizes with VGLUT2-containing vesicles in the EPN→LHb pathway, and Syt2 knockdown results in increased excitatory events within the LHb. Syt3 colocalizes with VGAT-containing vesicles in the EPN→LHb pathway, and Syt3 knockdown results in increased inhibitory events in the LHb. This distribution of calcium sensors provides a molecular explanation for how a single presynaptic terminal can independently tune excitatory and inhibitory output within the LHb. By segregating neurotransmitter-specific machinery into distinct vesicle pools—paired with either Syt2 for glutamate or Syt3 for GABA—the neuron effectively establishes a flexible “molecular switch”. Rather than being constrained by a fixed transmitter ratio, this organization allows the EPN→LHb pathway to dynamically shift signaling polarity in response to specific physiological needs.

### Shared mechanisms across dual-transmitter pathways

These findings expand on earlier observations of dual glutamate/GABA release from mesohabenular and pallidal afferents, which established that transmitter co-release in the LHb modulates mood-related behaviors^8,13^. The identification of Syt2 and Syt3 as transmitter-specific Ca^2+^ sensors refines this model by introducing a presynaptic mechanism for temporal and kinetic control. Syt2 is a low-affinity, fast-binding Ca^2+^ sensor associated with synchronous vesicle fusion^25^, whereas Syt3 exhibits higher Ca^2+^ affinity, slower kinetics, and governs asynchronous release^17,21,22,26^. The near-exclusive pairing of Syt2 with glutamatergic vesicles and Syt3 with GABAergic vesicles within the EPN→LHb pathway therefore offers a biophysical basis for differential release timing. Such temporal staggering could fine-tune postsynaptic integration within LHb neurons, generating dynamic excitatory–inhibitory balance rather than fixed transmitter ratios which may be a shared mechanism across dual release synapses. How can our observations be reconciled with a prior study^14^ reporting that EPN→LHb dual release terminals function via co-packaging of glutamate and GABA based on highly correlated trial-to-trial post-synaptic currents? The correlated signaling observed in the Kim et al. study can be re-interpreted as functional synchrony. While further studies are needed to fully reconcile these differences, the observation that both Syt2 and Syt3 isoforms are present within the same terminal implies that a single presynaptic action potential can trigger the parallel activation of both vesicle pools, mimicking the statistical signature of co-packaging.

### Synaptotagmin balance and behavioral regulation

Functionally, the proposed dual release mechanism may underlie the bidirectional behavioral effects attributed to the LHb in mood regulation. Pallido-habenular neurons influence both aversive and reward-related signaling, and manipulations that alter the glutamate/GABA release ratio modify depressive phenotypes in animal models^8,12,27^. Our data suggest that synaptotagmin isoform composition could act as a molecular switch controlling this ratio. Loss of Syt2 enhanced glutamatergic transmission, potentially increasing LHb excitability and promoting aversive output, whereas loss of Syt3 enhanced inhibitory tone. These complementary effects highlight a presynaptic mechanism by which Ca^2+^ sensor balance modulates circuit polarity and, ultimately, behavioral state.

Future work should address whether Syt2/Syt3 expression is dynamically regulated by activity, stress, or antidepressant treatment. Experience-dependent or neuromodulatory changes in isoform abundance could bias transmitter release toward excitation or inhibition, contributing to the plasticity of LHb output. At a broader level, the segregation of synaptotagmin isoforms to transmitter-specific vesicle pools may represent a general principle of dual-release organization throughout the brain. Extending these observations with high-resolution ultrastructural mapping, optogenetic stimulation, and behavioral assays will clarify how isoform diversity shapes computation in mixed-transmitter circuits. Together, our findings provide a mechanistic framework linking presynaptic Ca^2+^ sensor identity to the fine control of excitatory and inhibitory signaling in the habenula.

## MATERIALS AND METHODS

### Animals

Both C57BL/6J (RRID:IMSR_JAX:000664) and VGluT2-IRES:Cre (RRID:IMSR_JAX:016963) mice were purchased from The Jackson Laboratory (Bar Harbor, ME). Mice were maintained in a colony room with a 12-hr light/dark cycle with food and water access ad libitum. Both male and female mice between 4 and 6 months of age at the time of surgical procedures and were selected at random. All animal procedures were performed in accordance with the National Institutes of Health Guide for the Care and Use of Laboratory Animals and approved by the University of Colorado Boulder Institutional Animal Care and Use Committee (IACUC).

### Surgical procedures

Mice were rapidly anesthetized using 3% isoflurane and head-fixed in a Kopf stereotaxic instrument (Model 963), where anesthesia was maintained at 1–2% for the duration of surgery. Sedation depth was verified using toe pinch and leg extension reflex tests. Body temperature was maintained on an electric heating pad. The analgesic ER Buprenorphine (1 mg/kg, subcutaneous) was administered at the start of the procedure. The surgical site was prepared by applying Nair hair removal cream, which was removed using sterile cotton tips. The exposed skin surface was then sterilized with three alternating washes of betadine and 70% ethanol. A midline incision was made, and soft tissue was cleared from the skull with sterile cotton swabs. The head position was adjusted to achieve a Bregma–Lambda dorsal–ventral difference <0.05 mm. Craniotomies were drilled (0.5 mm burr) at stereotaxic coordinates relative to Bregma while avoiding damage to blood vessels or meninges. Injection volume (400 nL) and flow rate (2 nL/s) were controlled using a Nanoject III programmable nanoliter injector (Drummond Scientific, Broomall, PA). After injection, the needle remained in place for 9 min at the target depth and 1 min at +0.1 mm dorsal to minimize reflux. The incision was closed with 3M VetBond, and a bolus of sterile saline was administered subcutaneously for hydration. Animals were transferred to a recovery cage positioned half on a heating pad and monitored until fully ambulatory. Supplemental DietGel® and HydroGel® (ClearH2O, Westbrook, ME) were provided. Mice were singly housed until experimental use or tissue collection, typically 3–5 days post-injection.

### Intracranial Injection of AAV-hSyn-DIO-CoChR and FANA-ASO

Mice were anesthetized with 1-3% Isoflurane and secured in a stereotactic frame. AAV8-hSyn-FLEX-CoChR-GFP (UNC Vector Core), titre: 2-5 x 10^12 was injected into the EPN (AP: −1.2, ML: +−1.8, DV: −4.35 (females) −4.45 (males). Injection volume (400 nL) and flow rate (100 nL/min) were controlled with a microinjection pump (Micro4; World Precision Instruments, Sarasota, FL). Following injection, the needle was left in place for an additional 10 min for virus diffusion. Following the same procedures above, antisense oligonucleotides (FANA-ASOs, AUM Biotech) targeting either Syt2 or Syt3 murine transcripts (Syt2:NM_009307.3 and Syt3:NM_016663.3) were used to knockdown Syt2 or Syt3 in the EPN. Specifically, Syt2 was target with the following 5 ASOs: Syt2-ASO1 (21 nt) 5’-CATAAGGGTCTGATGTGCCAC-3’, Syt2-ASO2 (21nt) 5’-TAGATTGCCATCACCAGGGTC-3’, Syt2-ASO3 (21nt) 5’-TAGATTGCCATCACCAGGGTC-3’, Syt2-ASO3 (21nt) 5’-GTGTTCATGGGTACCTTCACC-3’, Syt2-ASO4 (21nt) 5’-AGTCTCTTACCGTTCTGCATC-3’, Syt2-ASO5 (21nt) 5’-AGGACTCGTTGAAGTAGGGGT-3’. Syt3 was targeted with the following 5 ASOs: Syt3-ASO1 (21 nt) 5’-ACACCTATCATTGGTATCAGC-3’, Syt3-ASO2 (21 nt) 5’-TCTGACCGATGAGCTTGGCTT-3’, Syt3-ASO3 (21nt) 5’-GTTCAACGTCTTCCTGTGCAC-3’, Syt3-ASO4 (20nt) 5’-TTCATTGTGGCCGATGCAGT-3’, Syt3-ASO5 (20nt), 5’-GATGTCTGGCTTGGTTTGAC-3’. The ASOs were injected into the EPN (coordinates from Bregma: AP −1.2 mm, ML ±1.8 mm, DV −4.35 mm [female] or −4.45 mm [male]) and the contralateral EPN was used as a control. For quantification of ASO spread, fluorescence control ASOs were injected at different concentrations, and the spread area was quantified by confocal microscopy (**Figure S1**).

### Perfusions

Mice were deeply anesthetized with an intraperitoneal overdose of Euthasol (200 mg/kg). Following loss of reflexes, transcardial perfusion was performed with 20 mL of ice-cold 1× PBS at a constant flow rate (approx. 2 mL/s), followed by 20 mL of ice-cold 4% paraformaldehyde (PFA). Brains were removed and post-fixed in 4% PFA for 24 h, then transferred to 30% sucrose for cryoprotection before sectioning.

### Fixed Sectioning

Brains were rinsed in 0.1 M phosphate buffer (PB) for ∼1 h before sectioning. The cerebellum and optic chiasm were removed, and the remaining tissue was secured to the stage of a Leica VT1000 vibratome using cyanoacrylate glue. Approximately 3.5 mm of tissue was trimmed before collecting 30 µm coronal slices (n = 100) spanning 6.3–7.8 mm. Cutting speed was set to 3.5 and frequency to 9.

### Immunohistochemistry (IHC)

All wash and staining steps were performed at 90 rpm on an orbital shaker. Fixed sections were washed three times in PBS, then blocked for 2 h at room temperature (RT). Primary antibody incubation occurred overnight at 4°C (1:500 dilution), followed by three PBS washes and 2 h secondary incubation at RT (1:1000 dilution).

### Microscopy

Sections were mounted in 0.1 M PB on glass slides and coverslipped with ProLong Diamond Antifade Mountant (Thermo Fisher). Slides were cured for 24 h at RT under light pressure and stored in the dark at 4°C. Imaging was performed using a Nikon A1 laser scanning confocal microscope. Localization images were collected at 4×, and analysis images at 40X or 100X (1024×1024 and 2048×2048 pixel resolution respectively). Images were analyzed using Fiji (RRID:SCR_002285) and Imaris (Bitplane: RRID:SCR_007370) for 3D reconstruction and colocalization analysis.

### Antibodies and Antibody Combinations

Antibodies were obtained from either Synaptic Systems or Jackson ImmunoResearch Laboratories and were aliquoted prior to use and a single aliquot was used for each experiment, **Table 1**.

**Table 1.**
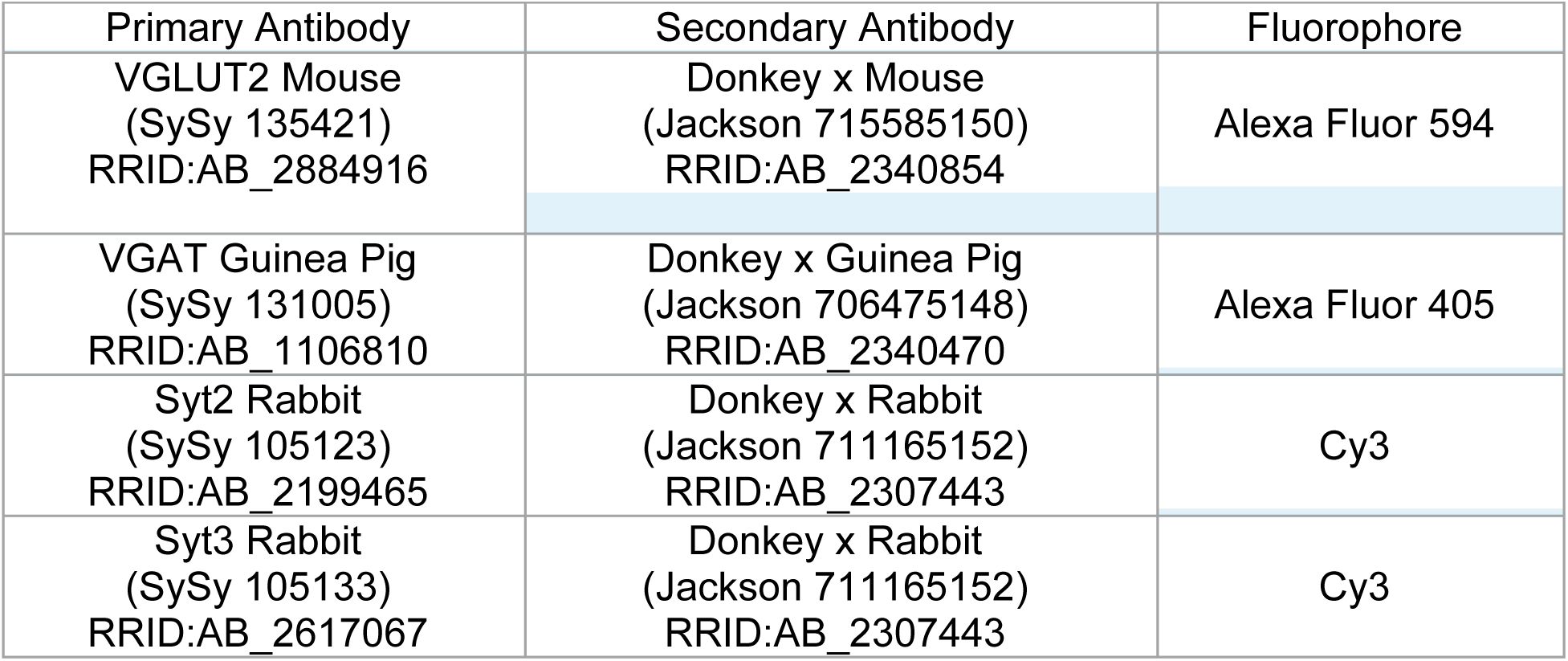
Primary and secondary antibodies and secondary fluorophore.

### Acute Brain Slices

Mice previously injected unilaterally with Syt2 or Syt3 ASOs as above were anesthetized with isoflurane and decapitated. Brains were immediately submerged in 4°C, carbogenated (95% O₂/5% CO₂) cutting ACSF containing (in mM): 110 sucrose, 5 glucose, 60 NaCl, 3 KCl, 1.25 NaH₂PO₄, 28 NaHCO₃, 0.6 ascorbate, 7 MgCl₂, 0.5 CaCl₂ (310 mOsm/kg, pH 7.3–7.4). Coronal slices (300 µm) containing the LHb were cut using a Leica VT1200S vibratome and incubated at 35°C for 30 min in standard ACSF (in mM: 25 glucose, 125 NaCl, 2.5 KCl, 1.25 NaH₂PO₄, 25 NaHCO₃, 1 MgCl₂, 2 CaCl₂; 310 mOsm/kg, pH 7.3–7.4). Slices were then maintained at RT for 30 min before recording.

### Electrophysiology

Slices were visualized using an Olympus upright microscope equipped with 4× (0.10 NA) and 40× (0.80 NA) objectives, IR-DIC optics, a CoolSNAP EZ camera (Photometrics), and Micro-Manager 1.4 software (Open Imaging). Whole-cell voltage-clamp recordings were obtained from neurons in the ventrolateral LHb, where entopeduncular (EP) inputs are densest. Recording pipettes (2–5 MΩ, borosilicate glass, 1.5 OD; Sutter Instrument) were filled with an internal solution containing (in mM): 120 CsMeSO₃, 10 HEPES, 0.5 EGTA, 8 NaCl, 10 Na-phosphocreatine, 1 QX-314, 4 MgATP, 0.4 Na₂GTP (pH 7.3–7.4, 290 mOsm/kg). D-AP5 (50 µM) and tetrodotoxin (TTX, 1 µM) were bath-applied for 5 min before recording miniature EPSCs and IPSCs simultaneously at −20 mV for 5 min. Signals were amplified using a MultiClamp 700B amplifier, digitized via Digidata 1550 (Molecular Devices), and sampled at 20 kHz. Data were notch-filtered at 60 Hz and low-pass filtered at 1 kHz. Access resistance (Rₐ) was monitored throughout; data were excluded if Rₐ > 25 MΩ or changed >20%.

### Electrophysiological Recording Analysis

Miniature EPSCs and IPSCs were analyzed using threshold-based event detection, with the root mean square (RMS) of baseline noise set to 3×. A baseline search period of 10 ms and an average baseline window of 5 ms were used for each event, which was fit to a biexponential curve. Event rise times (10–90%) and decay constants (37% amplitude) were interpolated at 200 kHz for precise kinetic analysis.

### Western Blot Analysis and ASO Knockdown Quantification

Quantification was performed by comparing the contralateral EPN of injected mice using western blot analysis for Syt2 and Syt3 using Synaptophysin for normalization. Following electrophysiological experiments, the dissected LHb region tissue was rinsed twice with ice cold PBS and lysed with 50ml RIPA buffer supplemented with complete protease inhibitors and 0.25% Triton X-100 on ice for 20 minutes on a rocking platform. Lysed cells were collected into microcentrifuge tube and spun for 20 minutes at 10,000xg at 4°C. Supernatant was collected and used for total protein quantification using BCA Protein Assay (Pierce, #23228). After normalizing all the cell lysates to the same total protein concentration, 5μg of lysates were mixed with SDS loading buffer and without boiling were separated by SDS-PAGE (Any kD gels, BioRad#456-9035, 150V/50mA/gel for 90 min) followed by a transfer to nitrocellulose membranes using wet transfer system (Mini Trans Blot Cell) at low intensity conditions (30V, 90mA, 16h, 4^0^C) in 1xSDS running buffer and 20% methanol. Transfer efficiency was determined by staining with PonceauS stain and subsequent block in TBS-Tween containing 5% non-fat dry milk for 1h at RT on a shaker followed by incubation with primary antibodies; Rabbit polyclonal Syt2 (SySy #105123, RRID:AB_2199465, 1:1000); Rabbit polyclonal Syt3 (SySy #105133, RRID:AB_2199465,1:1000); Guinea pig polyclonal Syp1 (SySy #101308, RRID:AB_2924959, 1:5000), Rabbit monoclonal GAPDH (Invitrogen #4A9L6, RRID: AB_2849138). The membrane was washed with 1xTBS-Tween 3 times for 7minutes, then incubated with secondary antibodies, Goat anti-Rabbit-IgG-HRP (ThermoFisher #31466, RRID:AB_10960844) and Goat anti-Guinea Pig IgG-HRP (Invitrogen #A18769, RRID:AB_2535546), at 1:10,000 for 1h at RT. The immunoreactive bands were visualized using a Clarity Max^TM^ Western ECL substrate (Bio-Rad #1705062) and by the Chemidoc^TM^ MP imaging System (Bio-Rad).

### Image and Data Analysis

Stacks used for 3D image reconstruction comprised ∼100 10nm slices totaling a Z-section measuring 1uM through the LHb. The resulting file was imported directly into Imaris Cellsens for analysis. GFP labeled projections from the EPN were isolated in their respective channel, reconstructed as a volume, and saved as an object. The same step was completed for each fluorescence channel, creating volumetrically representative objects for both VGLUT2/VGAT and Syt2/Syt3 puncta. Proximity analysis was performed using these objects, measuring from the centroids. A threshold approach was used for co-localization determination. To account for spectral shift and natural variations in focal lengths or staining specificity, only puncta within 100nm of GFP labelled EPN projections were included in analysis. This filtered out any anomalous signal from background or partial captures. From the remaining puncta, only signals from fluorophores within 1000nm of one another were considered. This was to ensure that the labeled proteins were likely contained within the same synaptic terminal, the average size of which is approximately 1μm^28^. We then measured the distances between all remaining puncta, excluding duplicates, and exported all data points (supplementary material) for subsequent spatial analysis.

## Acknowledgments

We gratefully acknowledge funding from The MCDB Neurodegenerative Disease Fund (to M.H.B.S.), NIH-5T32GM145437 (to D.N.W.) and NIH-5R01NS120496 (to M.H.B.S.).

## Author contributions

M.H.B.S, D.H.R. and D.N.W. designed the study. D.N.W. collected and analyzed confocal microscopy data. J.K.K. collected and analyzed the electrophysiological data. K.E.W. and D.J.M. performed stereotactic injections. T.B. and D.N.W. performed western blot analysis. D.H.R. provided reagents and laboratory space. C.A.H. provided laboratory space and equipment. Z.R.D. provided laboratory space and equipment. M.H.B.S and D.N.W. wrote the manuscript which was reviewed and edited by all authors. The authors declare no competing interests.

## Animal Ethics

All animal procedures were performed in accordance with the National Institutes of Health Guide for the Care and Use of Laboratory Animals and approved by the University of Colorado Boulder Institutional Animal Care and Use Committee (protocol 1106.02).

